# Label-Free, Real-Time, In Vivo Optical Biopsy with a Handheld Quantitative Phase Microscope

**DOI:** 10.1101/2025.09.17.675943

**Authors:** Zhe Guang, Srinidhi Bharadwaj, Zhaobin Zhang, Stewart Neill, Jeffrey J. Olson, Francisco E. Robles

**Affiliations:** Wallace H. Coulter Department of Biomedical Engineering, Georgia Institute of Technology and Emory University, Atlanta, GA 30332, USA; Department of Medical Engineering, California Institute of Technology, Pasadena, CA 91125, USA; Department of Neurosurgery, Emory University School of Medicine, Atlanta, Georgia 30322, USA; Winship Cancer Institute, Emory University, Atlanta, Georgia 30322, USA; Department of Pathology & Laboratory Medicine, Emory University School of Medicine, Atlanta, Georgia 30322, USA

## Abstract

In this work we develop and demonstrate the utility of a compact, handheld quantitative phase imaging microscope that enables label-free, in vivo optical imaging of bulk tissues with clear cellular and subcellular histological detail in real-time. The proposed device overcomes significant challenges in optical imaging for in vivo applications, particularly for clinical human use. The approach uses quantitative oblique back illumination microscopy (qOBM) to obtain quantitative phase information of opaque samples using epi-illumination. The compact handheld probe achieves 0.8 µm lateral resolution, 5 µm axial resolution, 300 µm X 300 µm field of view, and operates at 25Hz in a wide-field (non-scanning) configuration, enabling real-time imaging. The probe is also inexpensive and has no moving components, making it robust. The utility of the probe is demonstrated in (1) human skin in vivo, (2) brain tumor tissue ex vivo from a murine tumor model and from discarded human tissue from neurosurgery, and (3) in vivo using healthy brain tissue from a large animal model (swine), simulating neurosurgical conditions. Given the clear cellular and subcellular histological detail (i.e., “optical biopsy”) obtained in real-time, combined with the ease-of-use and low-cost of the system, the proposed device has significant implications for a broad range of clinical applications.

## 1. Introduction

Optical imaging technologies offer significant clinical value as they can provide real-time, non-destructive, in-situ assessment of tissues for a wide range of applications^1–8^. However, despite significant progress, high-resolution imaging with clear cellular and subcellular contrast for in-vivo applications remains a significant challenge, particularly for human clinical use. The challenges are multifold. For instance, for fluorescence microscopy, only a limited number of exogenous fluorophores are available for human use (e.g., fluorescein), and their specificity is relatively limited, with contrast restricted to only the labeled structures (e.g., vasculature)^9,10^. Moreover, even approved exogenous fluorophores can have severe health adversities when applied in humans, ranging from headache, paresthesias, seizures and vomiting to more severe complications such as epileptic crisis, grand mal epilepsy, opisthotonous and peripheral nerve palsy^11^. For these reasons, label-free imaging is particularly appealing for clinical applications. Autofluorescence (AF), for example, is an extremely useful label-free imaging technique, but it provides modest cellular contrast and lacks subcellular detail, particularly in vivo^12,13^. Plus, because most endogenous fluorophores’ excitation is in the ultraviolet (UV) and near-UV spectral range, AF has very limited penetration depth and can produce significant phototoxic effects^14^. Most often AF is applied for spectroscopic analysis of live tissue rather than as an imaging tool^12,13^. Optical coherence tomography (OCT) and reflectance confocal microscopy (RCM) are arguably one of the most successful technologies for label-free, high-resolution imaging in humans, but they also suffer from poor cellular and (more so) subcellular contrast^2,3,6,15–19^. Label-free multiphoton microscopy technologies (e.g., second harmonic generation, autofluorescence, and coherent Raman) are extremely promising, but they are currently not widely used clinically due to concerns with photodamage^20–25^. Moreover, nonlinear systems are bulky, expensive, and require significant expertise to operate and maintain.

In this work we develop a handheld device that achieves label-free, high-resolution imaging with clear cellular and subcellular contrast in real-time, thus overcoming significant limitations of existing optical imaging technologies. The device enables quantitative phase imaging (QPI) of opaque samples, using a novel technology termed quantitative oblique back illumination microscopy (qOBM)^26–31^. Indeed, QPI enables label-free measurements of biological samples with exquisite subcellular contrast and nanometer-scale sensitivity, providing unprecedented access to important histological and biophysical information of cells and tissues non-invasively^32,33^. However, conventional QPI methods are restricted to thin samples due to their reliance to a transmission-based optical geometry, which hinders their overall clinical utility, particularly for in-vivo applications. With the recent advent of qOBM, which operates in epi-mode, the same level of rich histological and biophysical insight is available in thick tissues in 3D. Using a benchtop qOBM microscope system, we have demonstrated qOBM’s ability to image (1) animal and human thick tissue samples ex vivo^34–36^, (2) organoids/spheroids^37,38^, and (3) complex systems such as cells inside bioreactors^39,40^, among other biological applications^41^. With the benchtop system, we have also demonstrated qOBM’s ability to quantitatively identify brain tumor margins in an animal model ex vivo and visualize both high- and low-grade human brain tumors ex vivo, including with virtual hematoxylin and eosin (H&E) staining^35^.

Here we miniaturize the qOBM system to achieve a compact handheld formfactor that can be applied in a wide range of applications. The device operates in real time (25 Hz), has no moving/scanning components, achieves high resolution (0.8 µm lateral and 5 µm axial) with a 300 µm X 300 µm field of view, and provides clear cellular and subcellular detail based on the tissues’ refractive index composition. The device is also simple, comprising three optical elements and uses illumination from low-cost light emitting diodes (LEDs). We demonstrate the utility of the probe in vivo and ex vivo, in human and animal tissues, including skin and brain. The unique ability of the device to provide clear cellular and subcellular histological detail—effectively an “optical biopsy”—in real-time with a compact handheld formfactor can be transformative for many clinical and biomedical applications.

## 2. Results

### 2.1 Probe design and construction

The handheld qOBM probe (Fig. 1a-b) comprises of three optical elements: a 0.8 NA GRIN lens with 2.3X magnification (GRINTECH GT-MO-080-032-ACR-VISNIR); a compact long working distance (12 mm) 20X, 0.4 NA microscope objective (Olympus LMPLFLN); and a 75mm tube lens doublet (Thorlabs, AC127-075-AB). Light is detected using a compact CMOS camera (Basler: acA1920-150um). This configuration achieves a pixel resolution of 0.25 μm (object plane)/pixel (see Methods section for more details). The probe is enclosed in a custom ergonomic aluminum enclosure that is ∼3 cm in diameter at its widest point, and 20 cm long (including the camera). For illumination, 720 nm LEDs (LuxeonStar SinkPAD-II) are coupled to PMMA 0.5 NA fibers. The fibers are threaded to the distal end of the probe and are arranged 90-degrees from one another around the GRIN lens (Fig. 1(a)). The four LEDs are sequentially turned on in synchronization with the CMOS camera acquisition, operating at 100 Hz, for an overall qOBM acquisition rate of 25 Hz. All captured data from the camera are processed and saved in real time by a home-built LabView program on a PC.

**Figure 1:**
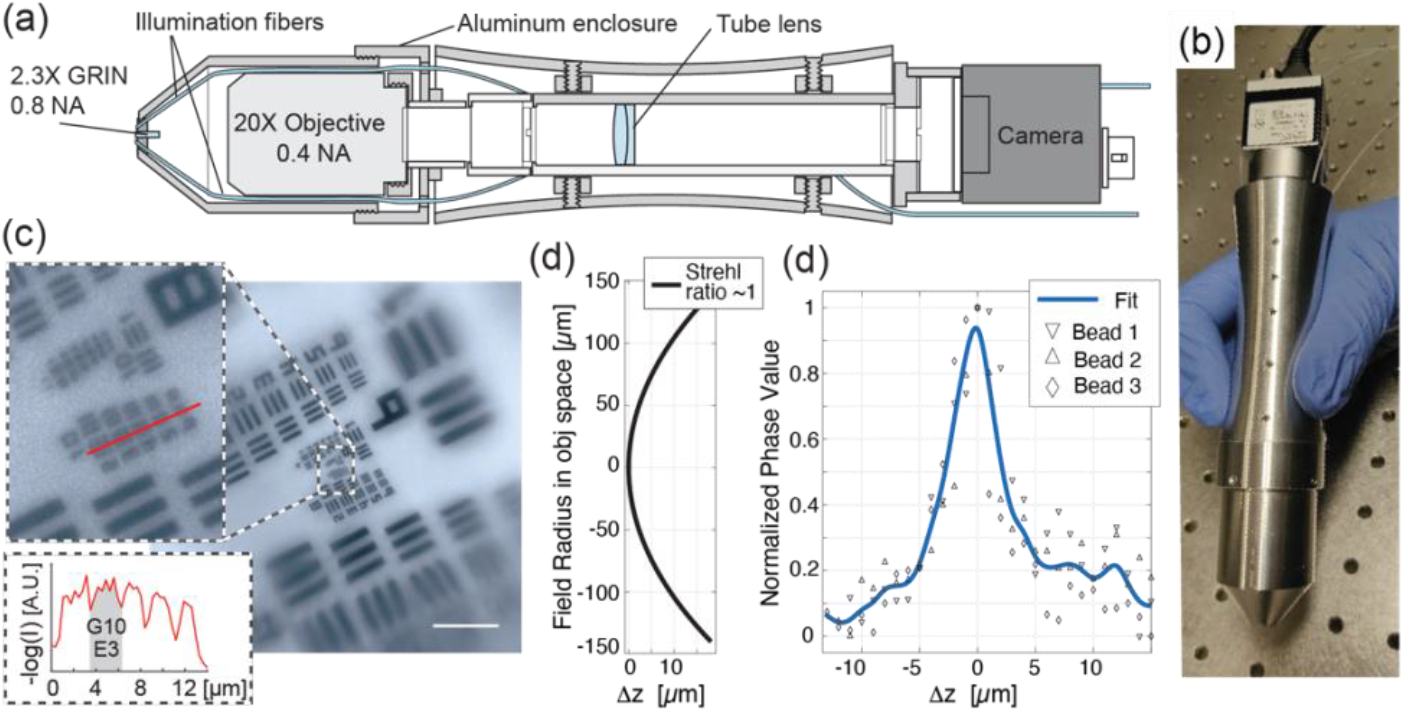
Handheld qOBM probe design. (a) Schematic and (b) photograph of the handheld probe. (c) Image of an USAF resolution test target acquired with the handheld probe (with scale bar 20 µm), and insets demonstrating a lateral resolution of ∼0.8 µm. (d) Field radius of system along which a Strahl ration of 1 is maintained (provided by manufacturer). (d) Axial resolution of the system is 5 µm, assessed experimentally using 1 µm beads.

As described in previous work, qOBM applies principles of oblique-back illumination to quantify phase in epi-mode and without a dedicated interferometer^26,27,31,42–44^. This process leverages multiple scattering within a thick sample (e.g., bulk tissue) to produce a virtual oblique light source within the thick medium. By principles of oblique illumination, combined with the low spatial coherence of the multiple-scattered light, cross-sectional phase information is obtained^26,27,42^. Using four separate image captures, two orthogonal differential phase contrast (DPC) images are obtained. Finally, to quantify the phase information, the two orthogonal DPC images are deconvolved in Fourier space using the system’s optical transfer function (OTF) and a computationally efficient Tikhonov deconvolution algorithm^26,27,43^. More details are provided in the Methods section.

Figure 1(c) shows a high-resolution test target image taken with the handheld qOBM system, where Group 10, Element 3 is resolvable, which corresponds with 1290 lp/mm or 0.78µm line spacing. This resolution of ∼0.8 µm is nearly diffraction limited at the center of the field of view (FOV). However, the probe, limited by the performance of the GRIN lens, suffers from significant field curvature aberrations^45^. As shown in Fig. 1(d), the field radius in the object space is 0.55 mm, which severely restricts the usable FOV when investigating thin, flat samples. However, for thick samples using a tomographic system, a near diffraction limited resolution is largely maintained along the curved field, thus enabling a much wider FOV with qOBM. As demonstrated in subsequent sections below, clear subcellular detail is observed along a FOV of 300 µm x 300 µm, with an axial focal shift of <20 µm from the center of the FOV to the periphery. Here the focus at the center of the FOV is set to ∼75 µm below the surface of the probe. Finally, the axial resolution of the handheld probe is 5 µm, experimentally assess using 1 µm polystyrene beads (Fig. 1(d)).

### 2.2 In-Vivo Human Skin Imaging

To first demonstrate the ability of handheld probe to provide histological grade information non-invasively, in vivo, and in real-time, we imaged the skin (forearm) of healthy volunteers. Figure 2 and Supplemental Videos 1-3 show representative results in which a number of interesting dermatological features can be seen, including clear cellular and subcellular detail from superficial structures (stratum corneum) through the epidermis and into the dermis where capillaries loops can be observed. Specifically, Fig. 2a clearly shows relatively large round structures (∼100 µm in diameter, green arrows) corresponding to the cross section of the rete ridges at the dermal-epidermal junction (DEJ), where we can observe basal cells and capillary loops with flowing blood cells. Figure 2b (orange arrow) shows large, high-refractive index structures corresponding to skin ridges largely composed of dead keratinocytes (effectively, folds in skin showing the stratum corneum). Figure 2c shows keratinocytes in the epidermis where structures at the center of the image are slightly deeper than those at the edges (due to field curvature) and likely correspond to the stratum spinosum and stratum granulosum, respectively. The two insets of Fig. 2c show keratinocytes of different sizes and shape as a result of being at slightly different depths along the epidermis. Figure 2d shows a different region of the epidermis where the upper inset shows a large cell, likely a Langerhans cell, surrounded by other (smaller) keratinocytes. The lower inset of Fig. 2d shows keratinocytes with a bright cytoplasm, likely indicating the presence of melanin. Finally, the upper inset of Fig. 2e shows red blood cells passing through a capillary loop in the upper dermis by the DEJ (Supplemental Video 3 provides a clearer view of blood flow). The lower inset of Fig. 2e shows basal cells which can be identified by their radial arrangement around the rete ridges. We also highlight that Supplemental Videos 1-2 clearly illustrates how smoothly the probe can be maneuvered to different regions of interest. Moreover, the video feedback can be viewed in real-time during acquisition.

**Figure 2.**
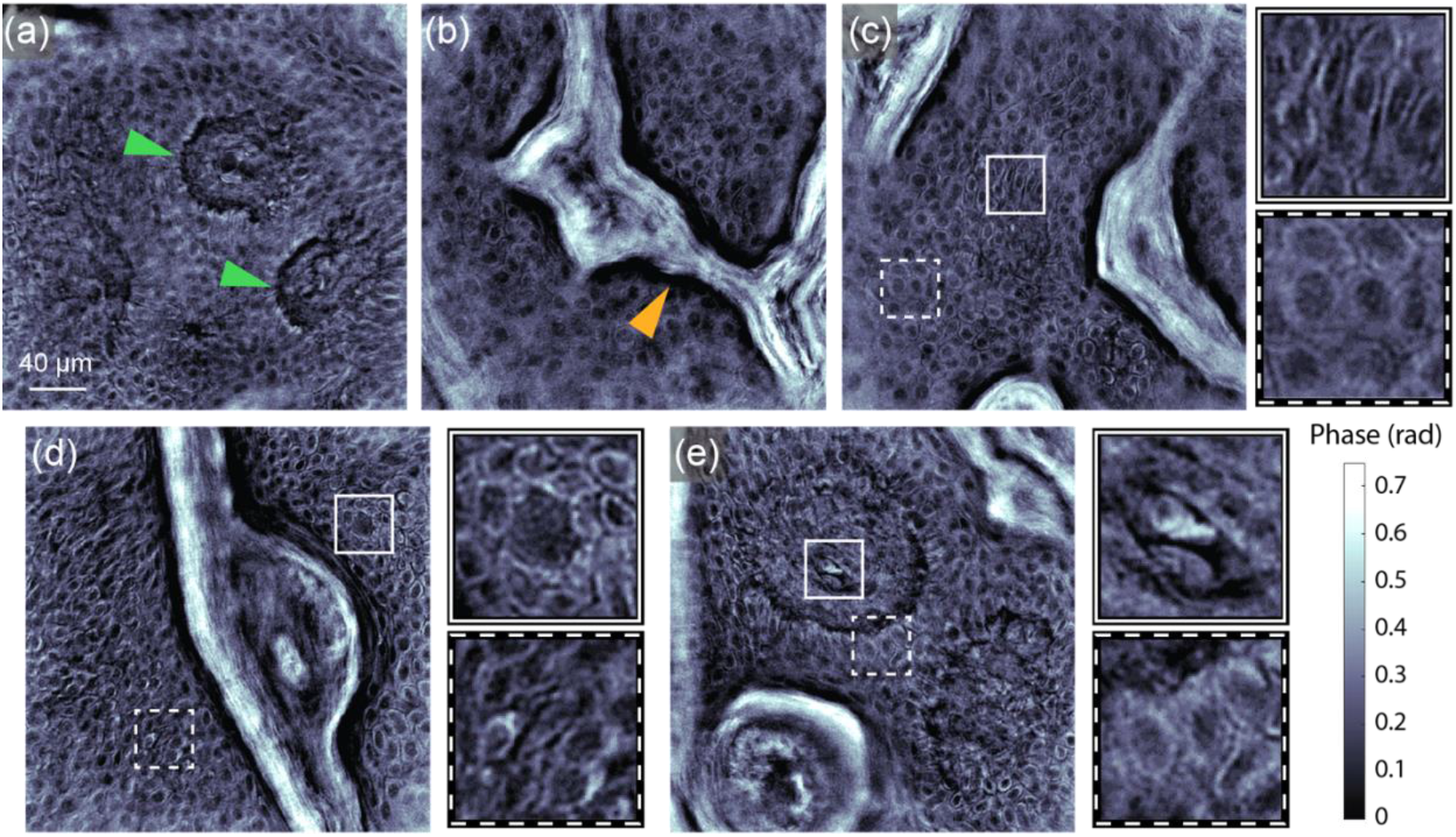
In-vivo human forearm skin images taken with the qOBM handheld probe. Green arrows highlight cross sections of the rete ridges at the dermal-epidermal junction (DEJ), where basal cells and capillary loops are observed. Orange arrow points to a skin fold illustrating the structure of the stratum corneum. Insets show various cells (keratinocytes and a larger Langerhans cell). Upper inset of (e) shows a red blood cell inside a capillary look in the DEJ. Scale bar and all insets are 40 µm. Supplemental Videos 1-3 illustrate real-time acquisition.

These results clearly demonstrate the ability of the qOBM handheld probe to provide unprecedented histological information (i.e., “optical biopsies”) of human tissue in vivo, in real-time without labels, using a simple, low-cost device. These capabilities are expected to have significant implications in dermatology.

### 2.3 Ex-Vivo Mouse Brain Imaging

Throughout the remainder of this work, we shift our focus to the use of the probe for applications related to surgical image guidance, specifically for neurosurgery. In this section, we use the handheld probe to image brain tissue ex vivo using healthy mice brain and a murine infiltrative tumor model of human gliomas (GL261). Healthy mice brain tissues were procured from unrelated experiments. Experiments using the GL261 glioblastoma murine model were approved the Emory IACUC (see methods sections for details). This model recapitulates the infiltrative nature of human gliomas and has been widely used to study aspects of glioma biology and therapeutic strategies^46,47^. All tissues were imaged within 1 hour of excision without fixation or staining—the handheld probe was simply placed in contact with the bulk tissue for imaging.

Figure 3 shows representative images acquired with the probe. Figures 3a-c correspond with brain tissue from healthy mice. Figure 3a, for example, shows a region of the cerebral cortex, with the neuropil—the network of interwoven neural and glial processes—clearly visible. The figure also shows a blood vessels with blood cells still inside, as seen by the upper inset. The lower inset of the figure shows a neuron, which in healthy tissue are typically relatively small and do not possess significant high phase (refractive index) contrast or prominent subnuclear structures. Figure 3b shows an area with many well-organized neurons in the hippocampus. As shown in the inset, small dots within the nuclei can be observed, which likely correspond with the nucleoli. Despite the high cellularity in this region, the well-organized structure, small round soma, and relatively low-contrast subnuclear content is indicative of healthy tissue^34,35,48^. Figure 3c shows another region of the cerebral cortex with different cell soma types present—likely, neurons with nucleoli also visible (upper inset) and glia (lower inset).

**Figure 3.**
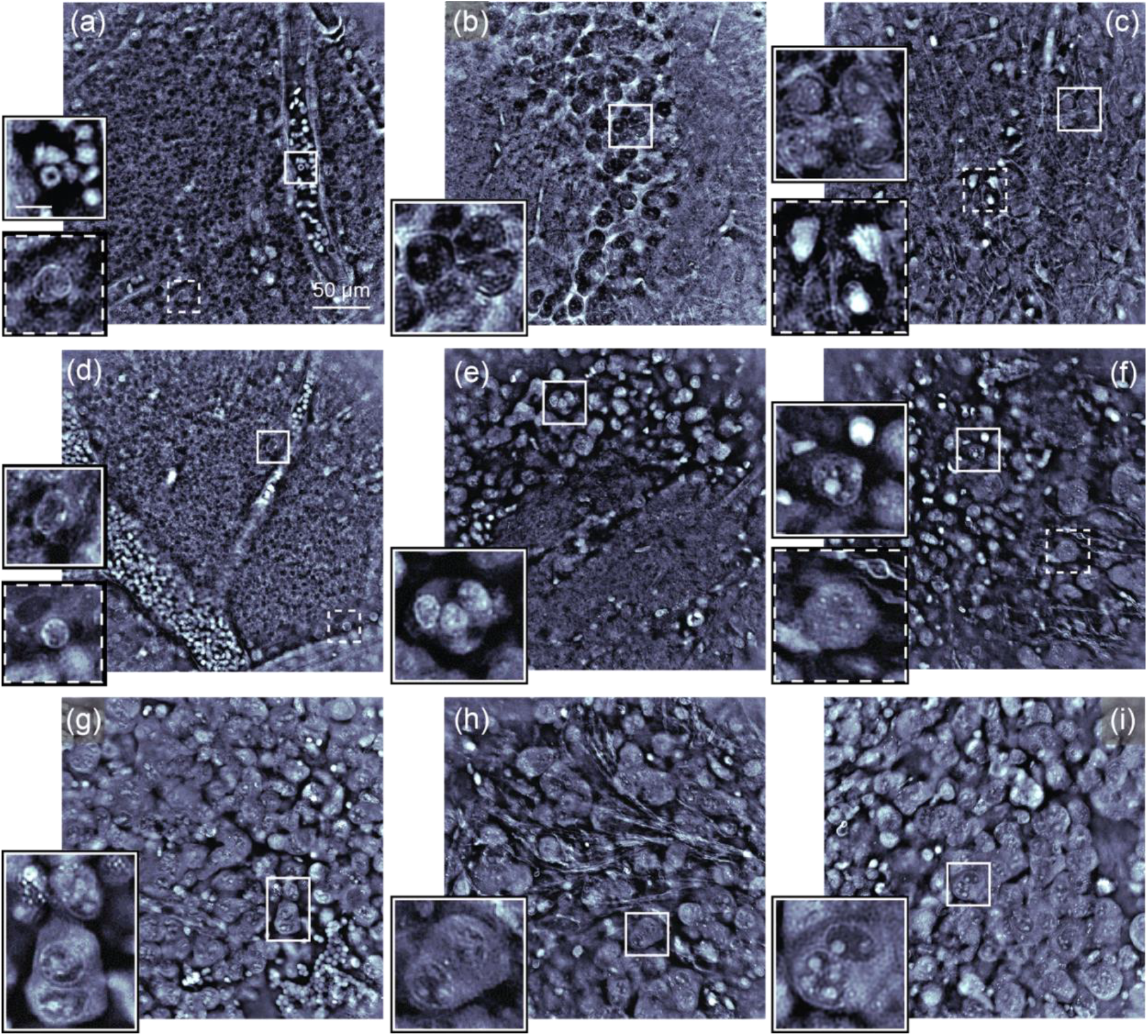
Ex-vivo fresh mouse brain images taken by the qOBM handheld probe. (a)-(c) Images of brain tissue from healthy mice. (d)-(i) Images of the brain tissue from the GL261 glioblastoma tumor model. (a) Normal neuropil with blood vessels with blood cells inside. (b) Hippocampal region. (c) Cortex region with various brain cells visible. (d) Healthy-appearing region from the GL261 model. (e) Infiltrative tumor margin. (f)-(i) Various bulk tumor regions exhibiting different morphologies. Insets in (g)-(i) are possible mitotic figures. The scale bar in the inset is 10 µm, and all insets have the same scale.

Figures 3d-i show representative images of the GL261 murine glioblastoma tumor model. Figure 3d shows a relatively normal region; here, a few different cell types can be seen near the blood vessels. The cell highlighted in the upper inset of Fig. 3d has similar appearance to the neuron-looking structure in Fig. 3a (lower inset), which suggest this could be a glial cell, such as an astrocyte. On the other hand, the smaller and rounder cell highlighted in the lower inset of 3d is more likely to be an immune cell (lymphocyte) that has accumulated with other similar-appearing cells near the tumor site. Figure 3e shows an infiltrative tumor margin, where the upper part shows bulk tumor tissue, and the lower part shows an admixture of normal-appearing neuropil and tumor cells, along with red blood cells and immune cells. Finally, Figs. 3f-i show various tumor regions expressing different morphologies. Note that the tumor cells are much larger, and their nuclei are clearly visible, enlarged, and contain numerous nucleoli (or chromocenters) which feature quite prominently and allow for ready identification of the tumor cells. These images also clearly show regions of high-density tumor cells and disruption of the normal morphology and architecture of the cortical and subcortical tissue. The insets of Fig. 3g-i show what appear to be several instances of multinucleation and nuclear atypia, which are pathological diagnostic indicators for malignant tumor cells. These results demonstrate the ability of the handheld probe to clearly visualize important cellular and subcellular structures related to brain tumor pathology.

### 2.4 Ex-Vivo Human Brain tumor Imaging

Next, we imaged human tissues discarded from neurosurgery. Bulk tissue specimens (fresh, unprocessed unless noted otherwise) were simply imaged with the qOBM handheld probe within 6 hrs of excision. Figures 4a-c show a glioblastoma WHO grade 4 sample with features consistent with a hypercellular glial neoplasm composed of cells with hyperchromatic nuclei. Similar to the H&E image, the qOBM images clearly show the sample hypercellularity as well as the hyperchromatic and pleomorphic nuclear features. In qOBM, hyperchromatic nuclei (indicated by a dark purple color in H&E) appear with a high phase (refractive index) value (i.e., bright nuclei) and sometimes with significant subnuclear structure. Pleomorphism refers to nuclei with irregular shapes and sizes, which is also clearly apparent in qOBM. Figures 4d-f show an example of a high-grade astrocytoma (WHO grade 3) which, as the H&E image shows, contains tumor cells with hyperchromatic nuclei arranged in a microcystic architecture. Again, these structures, particularly the cystic structure, are clearly evident in the qOBM phase images. It is important to emphasize that the qOBM images can be acquired in vivo intraoperatively and viewed in real-time (as shown below), while the H&E images require at best ∼20 mins for frozen sections (which can contain significant tissue artifacts) or many hours using more traditional Formalin-Fixed Paraffin-Embedded (FFPE) tissue sections (as shown here). The similarities in the structure between the gold-standard FFPE H&E images and qOBM images, underscores the ability of the handheld probe to provide label-free “optical biopsies” during neurosurgery in vivo.

**Figure 4.**
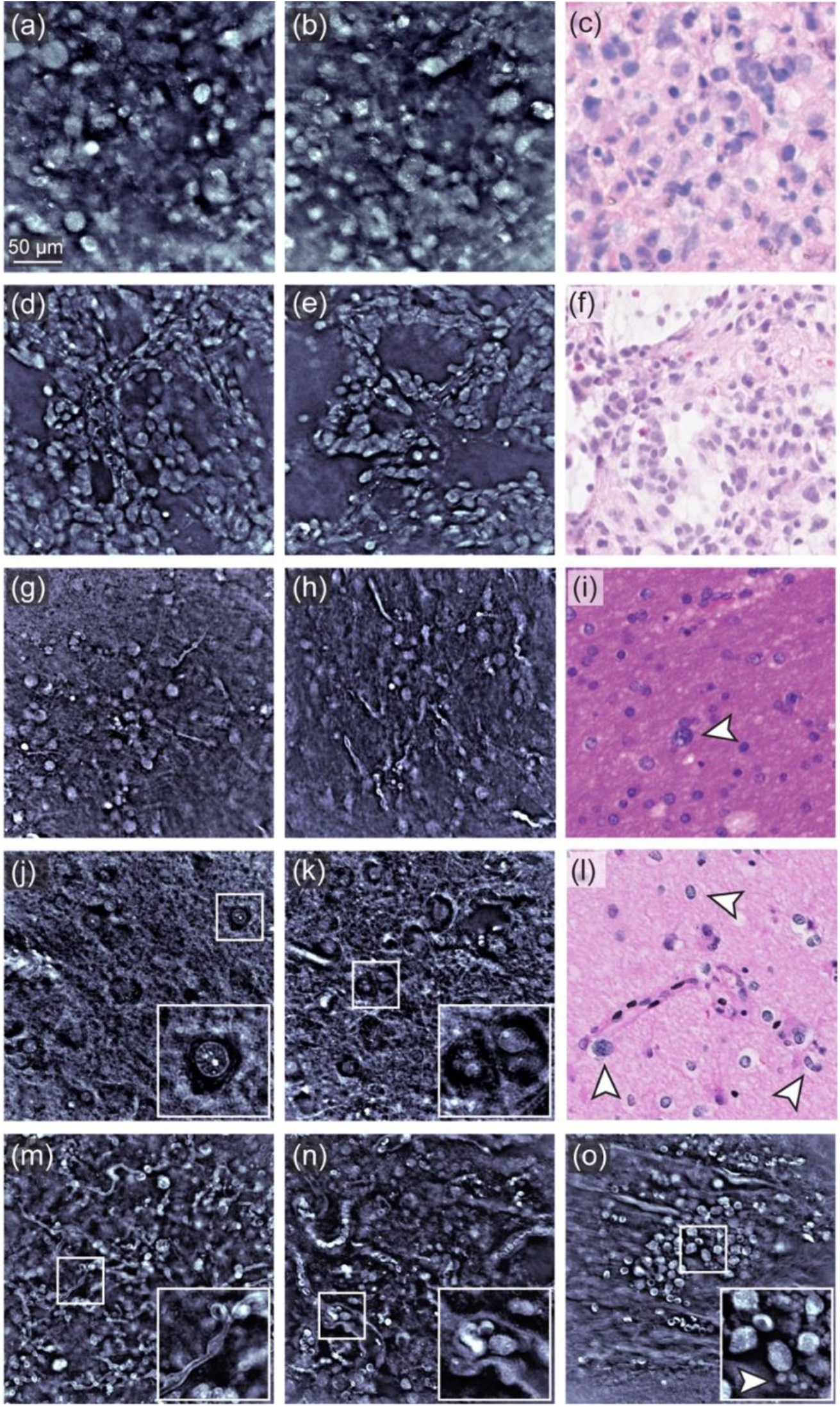
Images of ex-vivo bulk human tumors tissues discarded from neurosurgery taken with the qOBM handheld probe. (a)-(f) Phase images of two high-grade glioma acquired without tissue processing, with corresponding H&E image after tissue processing. (g)-(l) qOBM and H&E Images of two low-grade gliomas. qOBM probe images in (d)-(e) were acquired from bulk unprocessed tissue, while those in (j)-(k) were fixed but otherwise intact. (m)-(o) Show additional features visible in the handheld probe that are not readily observed under traditional histology, including myelinated axonal processes and small blood vessel with red and white blood cells inside, and platelets. Arrows in (i) and (l) show examples of tumor cells, as describe in the main text. Arrow in (o) highlights platelets. All insets are 50 µm x 50 µm.

Figures 4g-i show an example of a low-grade diffuse glioma, specifically a WHO grade 2 astrocytoma. This sample shows hyperchromatic, rounded to irregular tumor cells (see arrow in Fig. 4i). In the qOBM images, we can observe the increased nuclear density of the tumor and relatively high phase (refractive index) values of the nuclei, with some cells showing a granular subnuclear structure, consistent with the hyperchromatic appearance of the cells in H&E. Figures 4j-l show another low-grade (WHO grade 2) astrocytoma (this bulk tissue specimen was fixed prior to imaging but was otherwise intact). The cells here show round to irregular, hyperchromatic and focally pleomorphic nuclei, which are characteristic of this tumor type. This astrocytoma is less cellular than the previous example (Fig. 4g-i), which is evident in both the H&E and qOBM images. Despite the lower cellularity, the qOBM images in Fig. 4j-k show that the nuclei of this astrocytoma have substantial subnuclear structure (see inset of Fig 4j), which is consistent with the appearance of tumor cells. The inset of Fig. 4k also shows what appears to be large multinucleated tumor cells, which helps determine the presence of nuclear atypia and, in some tumor types, provides insight into the genetic subtype of the tumor^49^. It is important to highlight that diffuse low grade gliomas are notoriously difficult to identify, and currently there are no surgical tools available that can identify the margins of such tumors in vivo. The ability of the qOBM handheld probe to reveal nuclear hyperchromasia and nuclear substructures (which are difficult to assess on H&E, particularly frozen sections), further supports the potential utility of the device to identify tumor cells even in low-grade tumors and diffuse margins, which is critical for tumor margin evaluation. These results provide evidence that the qOBM handheld probe can be applied to address this critical unmet need in neurosurgery.

Figures 4m-o show additional structures found around low-grade diffuse tumors that are not observable even in traditional H&E histology. For example, Fig. 4m shows a region containing many tortuous myelinated axonal processes. These appear with high-contrast in the phase images due to the high-refractive index composition of the myelin, which contains lipids and proteins. These structures are typically dissolved during tissue processing and do not appear in traditional H&E histology. Figure 4n shows many small blood vessels again with a tortuous pattern. In many of the vessels blood cells can still be seen inside, and the inset shows a case where both red blood cells and a larger white blood cell can be seen. Figure 4o shows a site where a blood vessel had ruptured, and blood cells accumulated. Interestingly, white blood cells appear to be more abundant than even red blood cells, potentially as a result of the immune response to the tumor^50^. Small circular structures can also be observed (see arrow in the inset of Fig. 4o) which are likely platelets.

### 2.5 In-Vivo Large Animal Brain Imaging

As a final demonstration of the utility of the probe, we imaged the brain of a healthy Yorkshire swine in vivo. Experiments were approved by IACUC (T324P-TR). Typical neurosurgical conditions used in humans were replicated, including the craniotomy and cauterization to control bleeding (see methods section for more details). Moreover, the probe was encased in a sterile plastic sleeve for sterility, which would be required for human clinical use.

As Fig. 5 and Supplemental videos 4-6 show, the probe enables clear cellular and subcellular detail at video rate in vivo. The videos clearly show that the neurosurgeon (pictured in Fig. 5b) is able to smoothly maneuver the probe to desired locations in real-time while maintaining high image quality. Again, the probe provides real-time feedback (Fig. 5a). In the videos, pulsatile motion from the heartbeat can be observed; however, this does not produce imaging artifacts and does not result in motion blur of the surrounding tissue as engagement of the probe at the surface being imaged dampens that physiologic motion. Similarly, as observed in the videos, animal respiration does not affect image acquisition. The images (Fig. 5) show various regions of the cortex where the neuropil, blood flow, and neurons can be observed. Some areas, such as that seen in Fig. 5c, correspond with the delicate septated nature of the subarachnoid space. Figures 5d-h, on the other hand, correspond with cerebral cortical regions. These results clearly demonstrate that the probe can be used to image brain tissue in vivo under typical neurosurgical conditions, providing histological information in a non-destructive manner in real-time without having to remove the tissue (i.e., non-invasive, “optical biopsy”).

**Figure 5.**
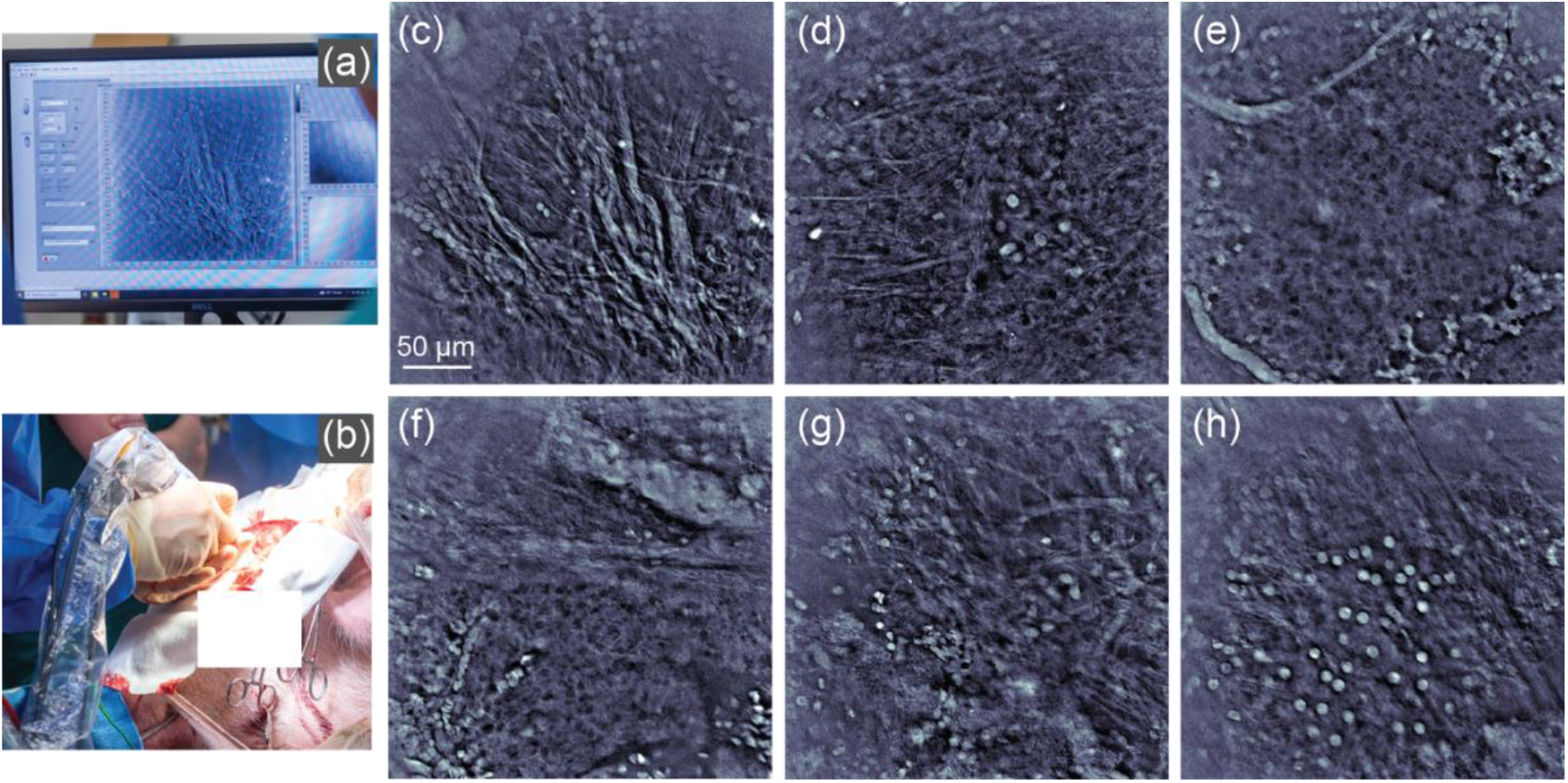
In-vivo, intraoperative qOBM brain imaging of a healthy large animal swine model. (a)-(b) Photographs taken during the surgical procedure, with (a) showing the real-time imaging feedback, and (b) showing the surgeon using the imaging probe. (c) qOBM image of subarachnoid space. (d)-(h) qOBM images of different cerebral cortical regions. Supplemental Videos 4-6 showcase the real-time imaging capabilities of the probe.

## Discussion and Conclusion

In this work, we have developed and demonstrated a novel handheld qOBM probe capable of label-free, high-resolution, non-invasive imaging with clear cellular and subcellular detail in real-time. These unique capabilities of the probe, together with its compact, robust, and cost-effective design, overcome significant limitations of existing optical imaging technologies, and make the device highly suitable for clinical in vivo applications. The level of subcellular histological detail provided, using a fast widefield imaging configuration, and without resorting to nonlinear optical phenomena, is unprecedented. Our demonstrations across human skin (in vivo), ex vivo murine brain tumors, discarded human glioma tissues, and a live swine brain model illustrate the broad applicability and potential clinical utility of the device. Its unique ability to deliver histology-grade images non-invasively and in real-time—effectively performing an optical biopsy—positions the handheld qOBM probe as a transformative tool for dermatological diagnostics, neurosurgical guidance, and many other clinical applications.

Future studies will focus on further clinical validation and exploration of additional medical applications, as well as incorporating additional functionalities. For example, the device in its current embodiment has a fixed focal plane, but simple modification can be made to enable volumetric 3D imaging. Similarly, the current operating wavelength is 720 nm, which enables a maximum penetration depth of ∼200 µm in brain tissue. In future work, longer wavelengths will be used to enable imaging at deeper depths, which will be particularly important when implementing 3D imaging capabilities. The device also has a limited field of view (300 µm x 300 µm), but live/iterative image stitching can be implemented in the future to enable acquisitions of large mosaics. Further efforts toward miniaturization will also be pursued to expand the versatility and clinical convenience of the device. Lastly, we have previously demonstrated successful image translation from qOBM phase images to virtual H&E histology in order to facilitate visual interpretation by pathologists^35^; we anticipate repeating this effort with the images acquired with the handheld probe, further improving the clinical utility of this device. With these future enhancements, we anticipate a significant expansion in the scope of clinical applications.

In conclusion, the handheld qOBM probe developed here holds great promise for improving clinical outcomes by enabling immediate/real-time, non-invasive, detailed histological evaluation directly at the point of care, potentially transforming patient management across various medical specialties. The development of this optical imaging device represents an important advancement towards accessible, accurate, and real-time diagnostics, which can streamline clinical workflows and ultimately contribute to better patient care and health outcomes.

## Methods

### Probe design

The probe has a 0.8 NA GRIN lens with 2.3X magnification (GRINTECH GT-MO-080-032-ACR-VISNIR) and a compact long working distance (12 mm) 20X, 0.4NA microscope objective (Olympus LMPLFLN). The image-side NA of the GRIN lens (0.32 NA) is well matched with that of the 20X objective (0.4NA), together providing the equivalent of a ∼50X, 0.8NA objective. This configuration yields a compact high-NA configuration with a long working distance from the microscope objective to the probe distal tip at a low-cost. Note that high NA (>0.7), high-magnification (>40X), long working distance (>10 mm) objectives are generally much bulkier and expensive in comparison. The long working distance from the microscope objective to the probe distal tip is important for two reasons: (1) it allows for a narrow, tapered probe tip, which helps users more precisely position the probe in a desired area, and (2) it allows for the illumination fibers to bend inside the probe aluminum housing from the objective to the probe distal tip without surpassing their bending radius (see Fig. 1(a)). From the objective, a ½” diameter, 75mm tube lens doublet (Thorlabs, AC127-075-AB) relays the light from the back aperture of the objective to a CMOS camera (Basler: acA1920-150um), providing a pixel resolution of 0.25 μm (object plane)/pixel.

### qOBM image processing

Two orthogonal DPC images are obtained by subtracting images acquired from the two sets of opposing illumination (I_1_ and I_2_), and the normalizing by their sum^43^, as described by Eq(1),

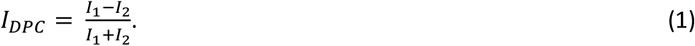

The two DPC images are linearly related to the phase (*ϕ*)^26,27^, as expressed in the Fourier domain in Eq.(2)

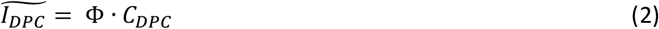

Where Φ is the Fourier transform of *ϕ*, and *C*_*DPC*_ is the optical transfer function (OTF) of the system. The OTF depends on the pupil function of the system, given by the NA of the objective, and the angular distribution of light at the focal plane. We have shown that the later term can be estimated numerically using photon transport simulations or measured experimentally^26–28^. To obtain quantitative phase information, the two orthogonal DPC images are deconvolved using the OTF of the system and a computationally efficient Tikhonov deconvolution^26,27^.

### Healthy Mice

Wild-type (WT) albino mice commonly used as control animals in neuroscience were used the study presented in section 2.3. The animals used here were donated from unrelated studies. Following IACUC approved protocols, mice were first anesthetized and then euthanized. Immediately following euthanasia, mouse brain tissues were dissected and imaged within 1 hour.

### GL261 glioblastoma model

Protocols were approved by the Emory IACUC. GL261 cells were cultured, harvested, and prepared as a suspension at 5×10^4^ cells/µL. Under sterile conditions, adult C57BL/6J mice were anesthetized using Isoflurane and then secured in a stereotaxic frame to inject the GL261 cells. Using a Hamilton syringe, a total of 1×10^5^ cells (2 µL) were stereotactically injected intracranially at a depth of 2.5–3.0 mm, followed by careful needle removal and sealing of the injection site. After the operation, animals receive analgesia, warmth, and close monitoring for neurological and behavioral changes. Brain and tumor tissues were harvested within 7–10 days.

### In vivo imaging of healthy Yorkshire swine

Experiments were approved by the IACUC (T324P-TR). The animal was previously used in an acute setting for surgical training. The swine craniotomy involved several steps. First, the swine was anesthetized, positioned prone in a stereotactic frame on a bed, and the surgical site on the head was shaved and sterilized. A midline incision was made along the scalp to expose the skull from the brow to the vertex, and small holes were drilled around the periphery of the proposed craniectomy. A side cutting drill was then used to connect the holes and create a bone flap. Having accomplished this, the bone flap was then carefully elevated to reveal the dura. The dura was incised to expose the frontal and parietal lobes on both sides to maximize opportunities to utilize the handheld probe in various orientations and configurations during the imaging trial.

## Supporting information

Video 1: Skin in vivo 1

Video 2: Skin in vivo 2

Video 3: Capillaries

Video 4: Brain in vivo 1

Video 5: Brain in vivo 2

Video 6: Brain in vivo 3

## Acknowledgments

We gratefully acknowledge assistance from Evan Golberg for facilitating experiments with the large swine animal model, and the Haider lab at Georgia Tech for providing discarded samples from healthy mice. We also acknowledge the following funding sources: Burroughs Wellcome Fund (CASI BWF 1014540); National Institute of General Medical Sciences (R35GM147437); the Georgia Research Alliance based in Atlanta, GA; Emory University Research Committee (URC); and Georgia Institute of Technology.

## References

1. Wilson, B. C. & Eu, D. Optical spectroscopy and imaging in surgical management of cancer patients. Transl. Biophotonics 4, (2022).

2. Rajadhyaksha, M., Marghoob, A., Rossi, A., Halpern, A. C. & Nehal, K. S. Reflectance confocal microscopy of skin in vivo: From bench to bedside. Lasers Surg. Med. 49, 7–19 (2017).

3. Kut, C. et al. Detection of human brain cancer infiltration ex vivo and in vivo using quantitative optical coherence tomography. Sci Transl Med 7, 292ra100–292ra100 (2015).

4. Richards-Kortum, R., Lorenzoni, C., Bagnato, V. S. & Schmeler, K. Optical imaging for screening and early cancer diagnosis in low-resource settings. Nat. Rev. Bioeng. 2, 25–43 (2024).

5. Hwang, K. et al. Handheld endomicroscope using a fiber-optic harmonograph enables real-time and in vivo confocal imaging of living cell morphology and capillary perfusion. Microsyst. Nanoeng. 6, 72 (2020).

6. Kulkarni, N. et al. Low-cost, chromatic confocal endomicroscope for cellular imaging in vivo. Biomed. Opt. Express 12, 5629 (2021).

7. Mueller, J. L. et al. An Accessible Laparoscope for Surgery in Low- and Middle-Income Countries. Ann Biomed Eng 49, 1657–1669 (2021).

8. Thompson, A. J. et al. The potential role of optical biopsy in the study and diagnosis of environmental enteric dysfunction. Nat. Rev. Gastroenterol. Hepatol. 14, 727–738 (2017).

9. Pogue, B. W. & Rosenthal, E. L. Review of successful pathways for regulatory approvals in open-field fluorescence-guided surgery. J. Biomed. Opt. 26, 030901–030901 (2021).

10. Seah, D., Cheng, Z. & Vendrell, M. Fluorescent Probes for Imaging in Humans: Where Are We Now? ACS Nano 17, 19478–19490 (2023).

11. Juneja, S. & Sandhu, K. Fluoroscein toxicity – Rare but dangerous. Indian J. Anaesth. 63, 674–675 (2019).

12. Croce, A. C. & Bottiroli, G. Autofluorescence spectroscopy and imaging: a tool for biomedical research and diagnosis. Eur. J. Histochem. 58, 2461 (2014).

13. Veld, D. C. G. D., Witjes, M. J. H., Sterenborg, H. J. C. M. & Roodenburg, J. L. N. The status of in vivo autofluorescence spectroscopy and imaging for oral oncology. Oral Oncol. 41, 117–131 (2005).

14. Clydesdale, G. J., Dandie, G. W. & Muller, H. K. Ultraviolet light induced injury: Immunological and inflammatory effects. Immunol. Cell Biol. 79, 547–568 (2001).

15. Bini, J. et al. Confocal mosaicing microscopy of human skin ex vivo: spectral analysis for digital staining to simulate histology-like appearance. J Biomed Opt 16, 076008-076008–8 (2011).

16. Dwyer, P. J., DiMarzio, C. A. & Rajadhyaksha, M. Confocal theta line-scanning microscope for imaging human tissues. Appl Optics 46, 1843 (2007).

17. Yin, C. et al. Label-free <italic>in vivo</italic> pathology of human epithelia with a high-speed handheld dual-axis confocal microscope. J Biomed Opt 24, 030501 (2019).

18. Gora, M. J., Suter, M. J., Tearney, G. J. & Li, X. Endoscopic optical coherence tomography: technologies and clinical applications [Invited]. Biomed. Opt. Express 8, 2405 (2017).

19. Liu, S. et al. Handheld optical coherence tomography for tissue imaging: current design and medical applications. Appl. Spectrosc. Rev. 60, 292–316 (2025).

20. Balu, M. et al. Distinguishing between Benign and Malignant Melanocytic Nevi by In Vivo Multiphoton Microscopy. Cancer Res 74, 2688–2697 (2014).

21. Balu, M. et al. In Vivo Multiphoton Microscopy of Basal Cell Carcinoma. Jama Dermatol 151, 1068–1074 (2015).

22. You, S. et al. Intravital imaging by simultaneous label-free autofluorescence-multiharmonic microscopy. Nat Commun 9, 2125 (2018).

23. Freudiger, C. W. et al. Multicolored stain-free histopathology with coherent Raman imaging. Lab Invest 92, 1492 (2012).

24. Hollon, T. C. et al. Near real-time intraoperative brain tumor diagnosis using stimulated Raman histology and deep neural networks. Nat Med 1–7 (2020) doi:10.1038/s41591-019-0715-9.

25. Ji, M. et al. Rapid, Label-Free Detection of Brain Tumors with Stimulated Raman Scattering Microscopy. Sci Transl Med 5, 201ra119 201ra119 (2013).

26. Ledwig, P. & Robles, F. E. Epi-mode tomographic quantitative phase imaging in thick scattering samples. Biomed Opt Express 10, 3605 (2019).

27. Ledwig, P. & Robles, F. E. Quantitative 3D refractive index tomography of opaque samples in epi-mode. Optica 8, 6 (2020).

28. Li, Z. et al. Experimental assessment of the optical transfer function for quantitative oblique back illumination microscopy (qOBM). Opt. Express 33, 5088 (2025).

29. Li, Z., Costa, P. C., Guang, Z., Filan, C. & Robles, F. E. GAN-based quantitative oblique back-illumination microscopy enables computationally efficient epi-mode refractive index tomography. Biomed. Opt. Express 15, 4764–4774 (2024).

30. Guang, Z., Ledwig, P., Costa, P. C., Filan, C. & Robles, F. E. Optimization of a flexible fiber-optic probe for epi-mode quantitative phase imaging. Opt Express 30, 17713 (2022).

31. Ledwig, P. & Robles, F. E. Partially coherent broadband 3D optical transfer functions with arbitrary temporal and angular power spectra. APL Photonics 8, 041301 (2023).

32. Park, Y., Depeursinge, C. & Popescu, G. Quantitative phase imaging in biomedicine. Nat Photonics 12, 1–12 (2018).

33. Huang, Z. & Cao, L. Quantitative phase imaging based on holography: trends and new perspectives. Light: Sci. Appl. 13, 145 (2024).

34. Costa, P. C. et al. Towards in-vivo label-free detection of brain tumor margins with epi-illumination tomographic quantitative phase imaging. Biomed Opt Express 12, 1621 (2021).

35. Abraham, T. M. et al. Label- and slide-free tissue histology using 3D epi-mode quantitative phase imaging and virtual hematoxylin and eosin staining. Optica 10, 1605 (2023).

36. Filan, C., Song, H., Platt, M. O. & Robles, F. E. Analysis of structural effects of sickle cell disease on brain vasculature of mice using three-dimensional quantitative phase imaging. J. Biomed. Opt. 28, 096501 (2023).

37. Serafini, C. E. et al. Non-invasive label-free imaging analysis pipeline for in situ characterization of 3D brain organoids. Sci. Rep. 14, 22331 (2024).

38. Serafini, C. E. et al. Longitudinal and continuous label-free monitoring of glioblastoma patient-derived tumor spheroid treatment response using quantitative oblique back illumination microscopy. Research Square preprint (2025) doi:10.21203/rs.3.rs-6529836/v.

39. Costa, P. C. et al. Functional imaging with dynamic quantitative oblique back-illumination microscopy. J Biomed Opt 27, 066502–066502 (2022).

40. Serafini, C. E. et al. Label-Free In-Line Characterization of Immune Cell Culture using Quantitative Phase Imaging. bioRxiv 2025.04.22.649403 (2025) doi:10.1101/2025.04.22.649403.

41. Serafini, C. E. et al. Label-free functional analysis of root-associated microbes with dynamic quantitative oblique back-illumination microscopy. Sci. Rep. 14, 5812 (2024).

42. Ford, T. N., Chu, K. K. & Mertz, J. Phase-gradient microscopy in thick tissue with oblique back-illumination. Nat Methods 9, 1195–1197 (2012).

43. Tian, L. & Waller, L. Quantitative differential phase contrast imaging in an LED array microscope. Opt Express 23, 11394–10 (2015).

44. Mehta, S. B. & Sheppard, C. J. R. Quantitative phase-gradient imaging at high resolution with asymmetric illumination-based differential phase contrast. Optics Letters 34, 1924–1926 (2009).

45. Matz, G., Messerschmidt, B. & Gross, H. Design and evaluation of new color-corrected rigid endomicroscopic high NA GRIN-objectives with a sub-micron resolution and large field of view. Opt. Express 24, 10987–11001 (2016).

46. Sahu, U., Barth, R. F., Otani, Y., McCormack, R. & Kaur, B. Rat and Mouse Brain Tumor Models for Experimental Neuro-Oncology Research. J. Neuropathol. Exp. Neurol. 81, 312–329 (2022).

47. Szatmári, T. et al. Detailed characterization of the mouse glioma 261 tumor model for experimental glioblastoma therapy. Cancer Sci 97, 546–553 (2006).

48. Su, Y. et al. Radix Rehmanniae Praeparata (Shu Dihuang) exerts neuroprotective effects on ICV-STZ-induced Alzheimer’s disease mice through modulation of INSR/IRS-1/AKT/GSK-3β signaling pathway and intestinal microbiota. Front. Pharmacol. 14, 1115387 (2023).

49. Mirzayans, R., Andrais, B. & Murray, D. Roles of Polyploid/Multinucleated Giant Cancer Cells in Metastasis and Disease Relapse Following Anticancer Treatment. Cancers 10, 118 (2018).

50. SenGupta, S., Hein, L. E. & Parent, C. A. The Recruitment of Neutrophils to the Tumor Microenvironment Is Regulated by Multiple Mediators. Front. Immunol. 12, 734188 (2021).

